# Evaluation of the sensitivity and specificity of three diagnostic tests for *Coxiella burnetii* infection in cattle and buffaloes in Punjab (India) using Bayesian latent class analysis

**DOI:** 10.1101/2021.07.23.453536

**Authors:** E. Meletis, R. Keshavamurthy, B.B. Singh, R.S. Aulakh, N.K. Dhand, P. Kostoulas

## Abstract

Q Fever is a zoonotic disease of significant animal and public health concern, caused by *Coxiella burnetii (C. burnetii)*, an obligate intracellular bacterium. This study was done to evaluate the sensitivity (*Se*) and specificity (*Sp*) of three diagnostic methods to diagnose *C. burnetii* infection in cattle and buffaloes in Punjab, India: an indirect ELISA method applied in serum samples and a trans-Polymerase Chain Reaction (trans-PCR) technique applied in milk samples and genital swabs. Bayesian Latent Class Models were developed following the STARD-BLCM reporting guidelines. Conditional independence was assumed between the tests, given (i) the different biological principle of ELISA and trans-PCR and (ii) the fact that the trans-PCR was performed on different tissues. The ELISA method in the serum samples showed the highest *Se* of 0.97 (95% Probability Intervals (PIs): 0.93; 0.99) compared to the trans-PCR method applied in milk samples 0.76 (0.62; 0.87) and genital swabs 0.7 (0.55; 0.82). The *Sps* of all tests were high, with trans-PCR in genital swabs recording the highest *Sp* of 0.99 (0.98; 1), while the *Sp* of trans-PCR in milk samples and ELISA in serum samples were 0.98 (0.96; 0.99) and 0.95 (0.93; 0.97) respectively. The study results show that none of the applied tests are perfect, therefore, a testing regimen based on the diagnostic characteristic of the tests may be considered for diagnosis of *C. burnetii*.

## 1. Introduction

Q Fever is a zoonotic disease that was first described by Edward Holbrook Derrick (Derrick, 1937) in Queensland. Q fever cases have been reported worldwide, except in New Zealand and Antarctica (Hilbink et al., 1993; Kaplan and Bertagna, 1955) and according to the OIE Terrestrial Animal Health Code, OIE countries and territories are obligated to report occurrences of the disease (OIE, 2018). Q stands for “query” and this designation was applied when the causative agent of the disease was unknown (OIE, 2018).

*Coxiella burnetii (C. burnetii)*, an obligate intracellular Gram-negative coccobacillary bacterium was identified as the causative agent of Q fever in 1938 (Raoult and Marrie, 1995; McDade, 1990). *C. burnetii*, as a Gram-negative bacterium can display two different phenotypes. Phase I bacteria are highly virulent, while Phase II bacteria are avirulent (Roest et al., 2013). *C. burnetii*, has been isolated from many domestic and wild animals, birds and arthropods; however, cattle, buffaloes, sheep, goats and humans are commonly affected (Babudieri, 1959). The species under investigation in this study are cattle and buffaloes. Further, many Ixodidae and Argasidae ticks are considered reservoirs of the bacterium (Mauren and Raoult, 1999). Even though prevalence studies of *C. burnetii* in ticks yield negligible estimates, bacterium transmission to tick species is considered feasible (Sprong et al., 2010).

Circulation of the bacterium has been described in wild animals’ populations and arthropods and dispersion of the bacterium in domestic populations can occur through air, direct contact and animal secretions/excretions (e.g., vaginal discharge, placenta, milk, feces, urine, saliva, amniotic fluid) (Angelakis and Raoult, 2010).

Limited information on the pathogenesis of *C. burnetii* in domestic animals is available, while under laboratory conditions several studies based on different animal models (e.g., guinea pigs, mice) have been conducted (Roest et al., 2013). In most cases, inhalation of aerosols or dust contaminated with birth fluids is described as the main route of infection (Waag, 2007). The pathogenesis and associated histopathological findings depend on the route of infection (Roest et al., 2013). Large *C. burnetii* concentrations are present in the infected placenta and amniotic fluid, while infected cows can shed the microorganism in milk for up to 32 months (Marrie, 2003). Guatteo et al. (2006) studied three different shedding routes - milk, vaginal mucus, feces - of the bacterium. Study results indicated no predominant *C. burnetii* shedding route and for the majority of shedder cows one shedding route was identified (Guatteo et al., 2006).

In livestock, the disease is usually subclinical, but the clinical form is associated with reproductive complications such as abortions, stillbirth, weak calves, repeat breeding and general clinical signs (e.g., anorexia) (Lang and Marie, 1990; EFSA, 2010; Keshavamurthy et al., 2019). Thus, *C. burnetii* excretion via feces, vaginal mucus and milk has been reported, sometimes independent of an abortion history (Roest et al., 2013).

Again, the host’s immune response to limit the infection has been studied in animal models. Specifically, during infection macrophages are the major target cells, while T-cells-associated with cellular immunity- and B-cells-associated with humoral immunity-, are critical for *C. burnetii* clearance after infection and tissue damage prevention, respectively. Antibody detection differs between the two Phases and the species (Roest et al., 2013).

Further, for the species under investigation in this study – cattle and buffaloes-the course of infection is considered similar (Lucchese et al., 2016).

In humans, the disease is observed in the (i) acute form, where flu-like symptoms, atypical pneumonia, hepatitis and cardiac involvement may be present and (ii) chronic form, that is more severe, than the acute, and fatal without appropriate therapy (Arricau-Bouvery and Rodolakis, 2005). Therefore, *C. burnetii* is characterized as a microorganism of great animal and public health concern.

Since there are no pathognomonic characteristics associated with *C. burnetii* infection, diagnosis poses a challenge. Many diagnostic tests, that are based on either the detection of the immune response of the host e.g., ELISA that detects antibodies directed against *C. burnetii* or the microorganism like Polymerase Chain Reaction (PCR) that detect bacterial DNA, have been used in epidemiological studies for *C. burnetii* to obtain estimates of the incidence and prevalence (Klemmer et al., 2018). However, prevalence estimation depends on the test’s sensitivity *(Se)* and specificity *(Sp)*, therefore, accurate diagnostic accuracy measures are important.

*C. burnetii* infection was first described in India in 1952 in a cattle herd and the first human case was reported in 1953 (Kalra and Taneja, 1954). Since then, the disease has been reported in several studies in India and an increasing trend in prevalence is reported, outlining *C. burnetii* as a potential threat to public health (Randhawa et al., 1973; Sodhi et. al., 1980; Vaidya et al., 2010; Keshavamurthy et. al., 2019; Shome et al., 2019; Yadav et al., 2019; Dhaka et al., 2020). Reported prevalence estimates from studies in India, conducted in a frequentist framework, assume that the applied diagnostic tests have perfect *Se* and *Sp* or imperfect, but known, measures of test accuracy. Further, a wide range of hosts, that *C. burnetii* is proven to affect, have been studied to better understand the microorganism’s epidemiology (Vaidya et al., 2010).

Objective of this study is the diagnostic evaluation of the applied tests to detect *C. burnetii* infection in cattle and buffalo animals in Punjab, a state in northern India. Since no gold standard is reported for *C. burnetii*, the study was conducted in a Bayesian framework, following the STARD-BLCM guidelines (Kostoulas et al., 2017). The STARD-BLCM checklist is available as a supplementary material (see S1 Appendix).

## 2. Materials and methods

### 2.1. Ethics approval

This study was approved by the Institutional Animal Ethical Committee, Guru Angad Dev Veterinary and Animal Sciences University (GADVASU/2017/IAEC/42/02).

### 2.2. Study design

The study design is presented in detail elsewhere (Keshavamurthy et al., 2019). Briefly, a multi-stage sampling design was performed. Twenty-two villages, one per district, of the state were selected. Further, the number of households sampled in each village was selected, proportional to the number of households in a village. Overall, 179 households (cattle or buffalo herds) participated in the study. We worked towards collecting samples from all the animals for each household. However, many farmers were reluctant to provide samples from some of their animals. We recorded a response rate of 72.5% at the household level and 53.4% at the animal level, respectively (Keshavamurthy et al., 2019).

The sampling unit in this study were cattle and buffaloes. In the analysis cattle and buffaloes were considered as one population, referred as domestic bovine population, because (i) both species are members of Bovidae family (Michelizzi et al., 2010) and (ii) the course of the *C. burnetii* infection is similar in both species (Lucchese et al., 2016). Under this setting, the target population of the study was the domestic bovine population in the Punjab state.

### 2.3. Sample collection

Blood and genital swab samples were collected from the selected cattle and buffaloes. In addition, milk samples, from lactating female animals, were collected. Blood samples were collected aseptically from the jugular vein. Puncture area was cleaned with 70% alcohol and venipuncture was done using a fresh needle. Approximately 5 ml of blood was withdrawn from each animal in a sterile vacutainer. For the molecular study, genital swabs were collected from both male and female animals using sterile cotton swabs. Vaginal swab samples from female animals were collected by carefully inserting the swab into the vaginal cavity about 10 cm through followed by gently rotating the swab. In males, preputial swabs were collected by swabbing the penile and preputial surface. For the collection of milk samples from the lactating animals, udder and teats were cleaned using germicidal teat dip and three to four streams of milk was discarded before sampling to minimize risk of sample contamination. About 15 ml of milk sample was collected using sterile screw-capped vials.

All samples were transported to the laboratory on the sampling day and stored at −20°C. The sera were separated within 24h in the sterile cryovials before storing at −20°C until screened (Keshavamurthy et al., 2019).

Overall, 610 blood samples, 610 genital swabs and 361 milk samples were collected from the study population. Therefore, the samples were split into two subpopulations; subpopulation-1 (subp_1) includes lactating female animals and subpopulation-2 (subp_2) includes male and non-lactating female animals. Overall, 400 of the sampled animals (65.5.%) were lactating and 221 (36.2%) were pregnant.

### 2.4. Laboratory tests

Milk samples and genital swabs were screened with PCR to detect bacterial DNA. Specifically, a trans-PCR assay was used to detect *C. burnetii* particles based on two transposon-like repetitive regions of the microorganism: Trans1: 5’-TAT GTA TCC ACC GTA GCC AGT C-3’ and Trans2: 5’-CCC AAC AAC ACC TCC TTA TTC-3’ (Willems et al., 1994).

The serum samples were screened for *C. burnetii* antibodies using the commercial Q Fever indirect ELISA kit with Phase I and Phase II (ELISA Kit for serodiagnosis of Q Fever in cattle and small ruminants, Monowell, Bio-X Diagnostics, Rochefort, Belgique). The two Phases capture both the acute and chronic infection form. In particular, IgG antibody titers against Phase I antigens are elevated during the acute phase, whereas IgG antibody titers against Phase I and Phase II are elevated during the chronic phase (ELISA Kit for serodiagnosis of Q Fever in cattle and small ruminants, Monowell, Bio-X Diagnostics, Rochefort, Belgique). ELISA quantifies the immune response of the host against *C. burnetii* and does not provide information about the presence or absence of the bacterium. Broadly, serological techniques are useful for screening purposes e.g., monitor the vaccination effectiveness. This is not applicable in India, because a vaccination program for *C. burnetii* does not exist (Dhaka et al., 2020).

### 2.5. Bayesian latent class analysis

The diagnostic accuracy of the applied tests was estimated using a Bayesian latent class model (BLCM). Traditionally, in the absence of a “gold standard”, latent class models (Hui and Walter, 1980) can be used to obtain unbiased estimates. Over the last decades, Bayesian framework has been applied in latent class analyses, due to their flexibility, incorporation of prior knowledge and software availability (Enøe et al., 2000; Branscum et al., 2005). To ensure transparency and extrapolation of the study results, the STARD-BLCM reporting guidelines for diagnostic accuracy studies that use BLCMs were followed (Kostoulas et al., 2017).

#### 2.5.1. Definition of infection status

Explicit description of the biological principle of each applied test is crucial towards the structure of any BLCM model. Latent variables are hidden, or unknown and probabilistic estimates can be made for them in conjunction with what the tests actually detect (Walter and Irwig, 1988). In this study, the applied PCR technique in the milk samples (PCR-Milk) and the genital swabs (PCR-Genital) detects bacterial DNA i.e., presence or absence of the *C. burnetii* microorganism and ELISA measures antibodies titers i.e., immune host response (IgG ELISA). Therefore, two distinct biological principles are defined implying different latent states of the sampling unit i.e., it is possible that presence of the bacteria in the host does not always trigger host immune response.

Estimations were based on the cross-classified results (Table 1) of the applied tests in the two subpopulations described above.

**Table 1.** Cross-classified results of IgG ELISA, PCR-Genital and PCR-Milk Lactating animals

|  |  | PCR-Genital<sup>1</sup> |  |  |  | Total |
| --- | --- | --- | --- | --- | --- | --- |
|  |  | Positive |  | Negative |  |  |
|  |  | PCR-Milk<sup>2</sup> |  | PCR-Milk |  |  |
|  |  | Positive | Negative | Positive | Negative |  |
| IgG ELISA<sup>3</sup> | Positive | 1 | 0 | 3 | 23 | 27 |
|  | Negative | 0 | 3 | 7 | 324 | 334 |
| Total |  | 1 | 3 | 10 | 347 | 361 |

Non-lactating animals
|  |  | PCR-Genital |  |  |  | Total |
| --- | --- | --- | --- | --- | --- | --- |
|  |  | Positive |  | Negative |  |  |
| IgG ELISA | Positive | 0 |  | 6 |  | 6 |
|  | Negative | 0 |  | 243 |  | 243 |
| Total |  | 0 |  | 249 |  | 249 |

Polymerase Chain Reaction (PCR) in genital swabs
<sup>2</sup>PCR-Milk: PCR in milk samples
<sup>3</sup>IgG ELISA: ELISA in serum samples

#### 2.5.2. Model assumptions

BLCM models, in the absence of a gold standard, for *Se* and *Sp* estimation can be constructed under different assumptions. An applied set of assumptions adopted by Hui and Walter model (two tests – two populations) state that (i) the population is divided into two or more subpopulations in which two or more tests are evaluated (ii) *Se* and *Sp* of each test remain constant across both species and both subpopulations and (iii) all tests are conditionally independent given infection status (Toft et al., 2005). According to the literature, previous Bayesian latent-class analyses for *C. burnetii* infection indicate that the *Se* and *Sp* of ELISA do not vary between species (Lucchese et al., 2016). Conditional independence can be assumed, on the basis that ELISA and PCR do not measure similar biological processes (Gardner et al., 2000). Also, PCR-Milk and PCR-Genital were applied to different organs, therefore, presence (absence) of the infectious agent to one organ does not imply presence (absence) to other organs. Even though, the existence of distinct difference of the true prevalence between the subpopulations is proven to influence the precision of the estimates; this is not applicable in this case, i.e., the two subpopulations have the same true prevalence.

#### 2.5.3. Model description

Bayesian modelling extracts the posterior probability given prior information and the likelihood function. The likelihood is computed through a statistical model for the observed data.

The models for the two subpopulations were structured assuming that the various test combinations follow the multinomial distribution. Specifically,

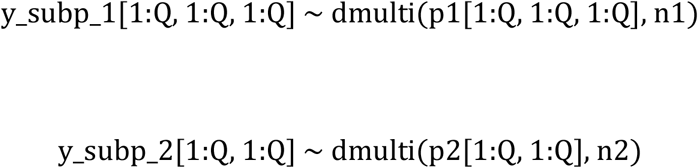

where y_subp_1 and y_subp_2 are the counts of various test combinations, Q = {1,2} the dichotomized test result i.e., 1 for positive and 2 for negative, n1 & n2 the two population sizes and p1 & p2 the probabilities of observing each test combination.

Based on this notation the frequencies of possible test outcomes can be calculated as followed:

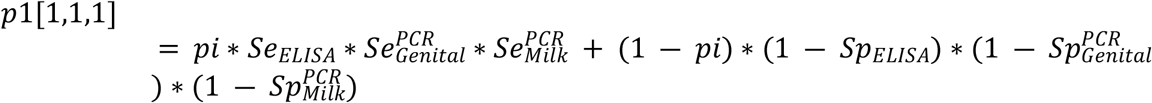

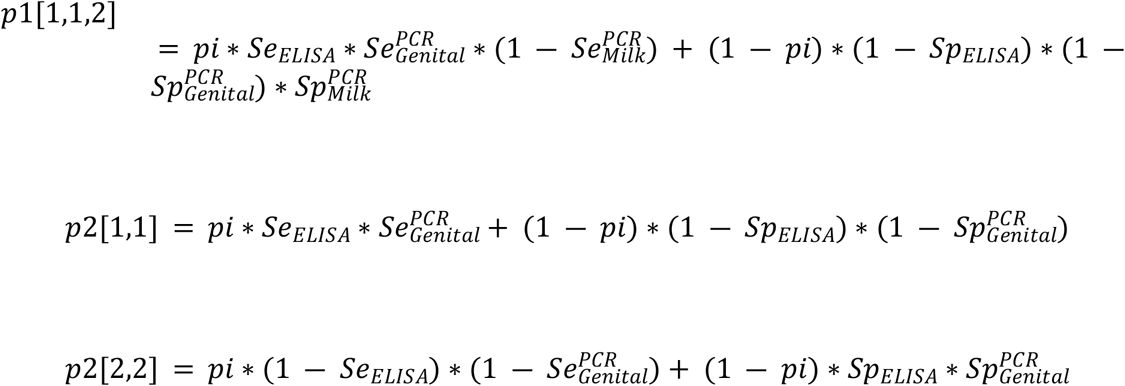

Under this setting, the parameters to be estimated are seven (i.e., Se_ELISA_, Se_PCR-Genital_, Se_PCR- Milk_, Sp_ELISA_, Sp_PCR-Genital_, Sp_PCR-Milk_ and pi (true prevalence)), while the degrees of freedom offered by the data are seven. Therefore, identifiability criteria are met and an uniform, noninformative, Beta prior distribution *Be (1,1)* can be adopted for all parameters of interest. However, degrees of freedom being higher than or equal to the number of parameters to be estimated is a necessary but not always sufficient condition to ensure identifiability. In this analysis, due to the sparsity of the observed data (i.e., zero cell observations for some of the cross-classified results – see Table 1) the ability of the model to estimate the associated parameters diminishes. Hence, informative priors were also introduced.

Prior information was supplied by one of the authors (B.B.S.), an epidemiologist, expert on zoonoses and co-leader of a national project on “Epidemiology, burden and control of zoonotic diseases in India”. Generally, not much is known about the differences in the *Se* and *Sp* of similar diagnostic tests in cattle and buffalo populations. A latent class analysis conducted using a Bayesian approach was performed to understand the differences in *Se* and *Sp* of ELISA in cattle and goats (Lucchese et al., 2016). The authors reported ELISA *Se* and *Sp* values to be 0.97 and 0.98 in cattle and 0.98, 0.83 in goats, respectively (Lucchese et al., 2016). In detail, estimates of the mean and the 95th percentile for all parameters were provided. Informative prior Beta distributions were calculated using the PriorGen R Package (Kostoulas, 2018) (Table 2).

**Table 2.** Mean and 95^th^ percentiles for the sensitivity (*Se*) and specificity (*Sp*) priors of PCR and ELISA and the corresponding Beta distributions, *Be(a, b)*. The prior information was provided by one of the co-authors (B.B.S.).

| Test | Parameter | Mean (%) | 95<sup>th</sup> percentile (%) | Be (a, b) |
| --- | --- | --- | --- | --- |
| PCR | *Se*<sup>*</sup> | 75 | 85 | *Be(32.58, 10.86)* |
|  | *Sp*<sup>**</sup> | 95 | 99 | *Be(40.78, 2.15)* |
| ELISA | *Se* | 97 | 99 | *Be(122.51, 3.79)* |
|  | *Sp* | 90 | 95 | *Be(11.74, 1.3)* |

#### 2.5.4. Markov Chain Monte Carlo convergence and software

Models were run in the freeware program OpenBUGS (Spiegelhalter et al., 2007). Parameter estimates were based on analytical summaries of 100,000 iterations of two chains after a burn-in phase of 5000 iterations. The tools described in Toft et al., 2007, were monitored to ensure occurrence of convergence. The OpenBUGS code for the final model is available as a supplementary material (see S2 Appendix).

#### 2.5.5. Sensitivity analysis

The influence of the data and the priors to the posterior estimates was examined by running the same model without informative prior values also. Different models under different set of assumptions were constructed to inspect the validity of the applied set of assumptions. Specifically, to ensure constant *Se* and *Sp* across species models to examine this assumption were structured.

Model selection was based on the DIC (Deviance Information Criterion) dialog box in OpenBUGS environment. The model with the smallest DIC is the model that best fits the data i.e., the model that would best predict a replicate dataset of the same structure as the currently observed (Spiegelhalter et al., 2002).

## 3. Results

The posterior medians and 95% probability intervals (PIs) for the *Se* and *Sp* of each diagnostic test are summarized in Table 3.

**Table 3.** Posterior medians and 95% probability intervals (PrIs) for the *Se* and *Sp* of each diagnostic test using informative Beta prior distributions for the *Se* and *Sp* of each diagnostic test described in Table 2.

| Test | Parameter | Posterior medians and 95% PrIs |
| --- | --- | --- |
| IgG ELISA | $Se$ | 0.97 (0.93; 0.99) |
| | $Sp$ | 0.95 (0.93; 0.97) |
| PCR-Genital | $Se$ | 0.7 (0.55; 0.82) |
| | $Sp$ | 0.99 (0.98; 1) |
| PCR-Milk | $Se$ | 0.76 (0.62; 0.87) |
| | $Sp$ | 0.98 (0.96; 0.99) |

IgG ELISA showed the highest *Se* with median 0.97 (95% PIs: 0.93; 0.99) compared to PCR-Milk 0.76 (0.62; 0.87) and PCR-genital 0.7 (0.55; 0.82). The *Sps* of all tests were high, with PCR-Genital recording the highest *Sp* median of 0.99 (0.98; 1), while the *Sp* of PCR-Milk and IgG ELISA were 0.98 (0.96; 0.99) and 0.95 (0.93; 0.97) respectively.

The acquired estimates without informative prior distributions are shown in S1 Table. Under this setting, the *Sp* estimates are similar in both scenarios. However, the *Se* estimates were lower; the *Se* of IgG ELISA was 0.63 (0.17; 0.98), PCR-Genital 0.18 (0.02; 0.77) and PCR-Milk was 0.6 (0.13; 0.98).

Sensitivity analysis was performed to validate the assumption of constant accuracy across both species. Parameter estimates produced by applying a model only to cattle were not substantially different, validating the assumption of constant accuracy across both species (S2 Table).

The model described in Section 2.5.3. was the one with the smallest DIC compared to the ones that were introduced in the Sensitivity Analysis section.

## 4. Discussion

In this study, BLCMs were used to estimate the Se and Sp of a trans-PCR applied in genital swabs and milk samples to detect *C. burnetii* DNA and an ELISA applied in serum samples that detects antibodies against *C. burnetii*, in cattle and buffaloes in Punjab, India. BLCMs account for the absence of a gold standard and allow parameter estimation merging two components (i) model structure, based on the observed data and (ii) incorporation of prior information (Kostoulas et al. 2017).

The study results show that all three tests are highly specific, with PCR-Genital yielding the higher Sp [0.99 (0.98; 1)], followed by PCR-Milk [0.98 (0.96; 0.99)] and IgG ELISA [0.95 (0.93; 0.99)]. On the other hand, IgG ELISA has the highest Se [0.97 (0.93; 0.99)], followed by PCR-Milk [0.76 (0.62; 0.87)] and PCR-Genital [0.7 (0.55; 0.82)]. This seems reasonable, as serological techniques are in general more sensitive than molecular diagnostic techniques, due to several reasons e.g., cross-reaction (Joseph et al., 1995). Overall, the posterior medians and 95% PIs for the PCR-Milk and PCR-Genital are comparable, indicating that both tests have the same diagnostic accuracy. Further, the Sps for these two tests, are not “prior-driven/dependent”, since both under informative and uniform, noninformative priors the posterior estimates are approximately the same. However, the reported estimates for the Ses of PCR-Milk, and especially PCR-Genital, seem to differ under noninformative and informative prior distributions [informative prior distributions; PCR-Milk 0.76 (0.62; 0.87) and PCR-Genital 0.7 (0.55; 0.82); noninformative prior distributions; PCR-Milk 0.6 (0.13; 0.98) and PCR-Genital 0.18 (0.02; 0.77)]. This is mainly due to the scarcity of the data i.e., small number of animals both positive to PCR-Milk and PCR-Genital. Further, the reported medians for the Se of PCR-Milk and PCR-Genital, using informative prior distributions, are included in the 95% PIs for the Ses under uniform, noninformative prior distributions. Implementation of informative prior distributions allows shrinkage of the 95% PI for the Ses. Therefore, the final Se estimates for the two PCRs can be considered reliable. As far as IgG-ELISA, even though, a high Sp is recorded, the Se seems to be “prior-driven/dependent”. However, the information provided by the data may not be enough, since only thirty-three animals were tested positive in IgG-ELISA [1 PCR-Milk+, PCR-Genital+, 3 PCR Milk+, PCR-Genital-, 23 PCR-Milk-, PCR-Genital-, 6 PCR-Genital-]. Therefore, IgG-ELISA Se posterior estimates using uniform, noninformative prior distribution cannot be considered reliable. Again, the median for the Se of IgG-ELISA, using informative priors is included in the 95% PI for the Se under a uniform, noninformative prior setting.

Studies on validation of diagnostic tests for *C. burnetii* infection using BLCMs have been conducted in cattle, goats, sheep etc. (Luchesse et al., 2016; Paul et al., 2013). The reported Se and Sp of the tests in our study are comparable between studies and similar between different species e.g., sheep and goats. (Abiri et al., 2019). The posterior medians and 95% PIs for the diagnostic characteristics of IgG-ELISA are similar to those reported in the literature and comparable with the estimates provided by the commercial ELISA kits manufacturers used for detection of antibodies against *C. burnetii* in serum samples from cattle (Luchese et al., 2016; Serrano-Pérez et al., 2015). The PCR method applied in milk samples has been evaluated in cattle (Nusinovici et al., 2015), in a Bayesian framework. The results from our study are similar with the ones reported in Nusinovici et al. (2015). The PCR method applied in genital swabs in cattle has not been evaluated; instead, PCR has been used for bacterial DNA detection in the farm environment (Nusinovici et al., 2015). On the other hand, PCR-Genital and PCR-Milk has been described in the sheep and goats (Abiri et al., 2019). The Sp for both PCRs and for the Se of PCR-Genital are similar, while the reported median and 95% PI for the Se of PCR-Milk in Abiri et al. (2019) is lower [0.42 (0.32; 0.59)].

Conditional independence between PCR-Milk and PCR-ELISA was assumed, since, the method was applied to different organs, even though it is based on the same biological principle. Further, primary shedding route has not been identified for *C. burnetii* and isolation of the bacterium from more than two organs is rare (Table 1) (Guatteo et al., 2006), i.e., presence (absence) of the infectious agent in the genital tract does not imply presence (absence) to milk. The conditional independence assumption for a PCR method applied in milk and vaginal secretions in sheep and goat samples to detect *C. burnetii* has been adopted (Abiri et al., 2019). Therefore, this assumption can be considered valid. Moreover, the specified model has seven degrees of freedom and seven parameters of interest. If conditional independence was not assumed, then two extra parameters for the covariance terms will be added, and the identifiability criteria will not be met (degrees of freedom higher than or equal to the number of parameters of interest), hence, our model would not converge. Therefore, introducing two covariance terms, accounting for conditional dependence between PCR-Milk & PCR-Genital would result in an unidentifiable model. Even though, in our case we adopt the results using informative prior distributions, we do so, due to the scarcity of the data (zero cell observations for animals positive to IgG-ELISA, PCR-Milk, PCR-Genital). Thus, informative prior distributions are added, instead of noninformative, to overcome the sparsity of the data. As shown in S2 Table the specified model using uniform, noninformative priors converges, but results to 95% PIs with high width.

Furthermore, the course of infection in the two species under investigation was considered similar, hence the diagnostic tests properties were assumed constant across species. Applying the model only to the cattle population yields posterior estimates similar to the ones after applying the final selected model (S2 Table). In India, risk factor investigation studies present contrasting results in the risk of occurrence of *C. burnetii* infection in bovine populations. A recent study conducted in Bihar and Assam states of India reported higher risk of infection for buffaloes than in cattle (28.0% compared to 13.6%, p=0.042) at the species level (Shome et al., 2019). However, only 25 buffaloes as compared to 719 cattle were included in this study (Shome et al, 2019). On the other hand, similar investigations in Punjab reported that cattle (adjusted Odds Ratio 3.37, 95% Confidence interval 1.23-9.20, p=0.02) were associated with larger odds of *C. burnetii* positive animal status than buffaloes (Keshavamurthy et al., 2020). Based on these results, our assumption that the course of infection and disease occurrence does not vary much in cattle and buffaloes at the species level is valid and there are other factors that need further investigation.

In this analysis, usage of informative priors improved the fit of the model, as indicated by the DIC. This is essential to account for the sparsity of the observed data. Assuming noninformative priors generates estimates with very wide PIs that do not allow safe conclusions about the tests’ accuracy.

## 5. Conclusion

This study was conducted to estimate the *Se* and *Sp* of three tests used for *C. burnetii* detection in Punjab, India. IgG ELISA achieved the highest *Se*, while PCR-Genital had the highest *Sp*. Using BLCMs, none of the applied tests showed perfect *Se* and *Sp*, and therefore, could not be used alone to diagnose *C. burnetii* infection in domestic bovine populations. Further, it is proven that the diagnostic accuracy of the tests does not vary between the two bovine species. In conclusion, better information on *C. burnetii* infection can be provided using a combination of diagnostic tests based on different biological principles i.e., detection of bacterial DNA and immune host response.

## Acknowledgments

The India Council for Agricultural Research, Government of India, New Delhi through the “Outreach Programme for Zoonotic Investigations” are thanked for their financial support of the present study.

## Declarations of interest

None

## Supporting information

**S1 Table. S1_Table_file**. Posterior medians and 95% PrIs for the sensitivity (Se) and specificity (Sp) of each diagnostic test using noninformative Beta prior distributions for all parameters of interest.

**S2 Table. S2_Table_file**. Posterior medians and 95% PrIs for the Se and Sp of each diagnostic test applying the model only to cattle population, assuming (i) informative Beta prior distributions for the Se and Sp of each diagnostic test and (ii) noninformative Beta prior distributions for all parameters of interest.

**S1 Appendix. STARD-BLCM Checklist**.

**S2 Appendix. OpenBUGS Code**

## Notes

### Competing Interest Statement

The authors have declared no competing interest.

## References

Abiri Z, Khalili M, Kostoulas P, Sharifi H, Rad M, Babaei H. Bayesian estimation of sensitivity and specificity of a PCR method to detect Coxiella burnetii in milk and vaginal secretions in sheep and goat samples. J. Dairy Sci. 2019;102: 4954–4959.

Angelakis E, Raoult D. Q fever. Vet. Mic. 2010;140: 297–309.

Arricau-Bouvery N, Rodolakis A. Is Q fever an emerging or re-emerging zoonosis?. Vet. Res. 2005;36: 327–349.

Babudieri B. Q fever: a zoonosis. Adv. Vet. Sci. 1959;5: 81

Branscum AJ, Gardner IA, Johnson WO. Estimation of diagnostic-test sensitivity and specificity through Bayesian modeling. Pre. Vet. Med. 2005;68: 145–163.

Burnet FM, Freeman M. Experimental Studies on the Virus of “Q” Fever. Med. J. Aus. 1937;2: 299–305.

Derrick EH. “Q” Fever, a New Fever Entity: Clinical Features, Diagnosis and Laboratory Investigation. Med. J. Aus. 1937;2: 281–299.

Dhaka P, Malik SVS, Yadav JP, Kumar M, Barbuddhe SB, Rawool DB. Apparent prevalence and risk factors of coxiellosis (Q fever) among dairy herds in India. PloS One, 2020;15: e0239260.

EFSA Panel on Animal Health and Welfare (AHAW). Scientific Opinion on Q fever. EFSA Journal. 2010;8: 1595.

Enøe C, Georgiadis MP, Johnson WO. Estimation of sensitivity and specificity of diagnostic tests and disease prevalence when the true disease state is unknown. Pre. Vet. Med. 2000;45: 61–81.

Gardner IA, Stryhn H, Lind P, Collins MT. Conditional dependence between tests affects the diagnosis and surveillance of animal diseases. Prev. Vet. Med. 2000;45: 107–122.

Guatteo R, Beaudeau F, Berri M, Rodolakis A, Joly A, Seegers H. Shedding routes of Coxiella burnetii in dairy cows: implications for detection and control. Vet. Res. 2006;37: 827–833.

Hilbink F, Penrose M, Kovacova E, Kazar J.. Q fever is absent from New Zealand. Int. J. Epidemiol. 1993;22: 945–949.

Hui, S.L., Walter, S.D., 1980. Estimating the error rates of diagnostic tests. Biometrics, 167–171.

Joseph L, Gyorkos TW, Coupal L. Bayesian estimation of disease prevalence and the parameters of diagnostic tests in the absence of a gold standard. Am. J. Epidemiol. 1995;141: 263–272.

Kalra SL, Taneja BL. Q fever in India: a serological survey. Indian J. Med. Res. 1954;42: 315–318.

Kaplan MM, Bertagna P. The geographical distribution of Q fever. Bull. World Health Organ. 1955;13: 829.

Keshavamurthy R, Singh BB, Kalambhe DG, Aulakh RS, Dhand NK. Prevalence of Coxiella burnetii in cattle and buffalo populations in Punjab, India. Prev. Vet. Med. 2019;166: 16–20.

Keshavamurthy R, Singh BB, Kalambhe, DG, Aulakh RS, Dhand NK. Identification of risk factors associated with Coxiella burnetii infection in cattle and buffaloes in India. Prev. Vet. Med. 2020;181: 105081.

Klemmer J, Njeru J, Emam A, El-Sayed A, Moawad AA, Henning K, et al. Q fever in Egypt: Epidemiological survey of Coxiella burnetii specific antibodies in cattle, buffaloes, sheep, goats and camels. PloS One. 2018;13: e0192188.

Kostoulas P. PriorGen: Generates Prior Distributions for Proportions. R 364 Packag. version, 2018;1: 365.

Kostoulas P, Nielsen SS, Branscum AJ, Johnson WO, Dendukuri N, Dhand NK, et al. STARD-BLCM: Standards for the Reporting of Diagnostic accuracy studies that use Bayesian Latent Class Models. Prev. Vet. Med. 2017;138: 37–47.

Lang GH, Marrie TJ. Coxiellosis (Q fever) in animals. In Q Fever: the disease. CRC Press, New York, USA; 1990. pp. 23–48.

Lucchese L, Capello K, Barberio A, Ceglie L, Guerrini E, Zuliani F, et al. Evaluation of serological tests for Q fever in ruminants using the latent class analysis. Clin. Res. Infect. Dis. 2016;3: 1–5.

Marrie TJ. Coxiella burnetii pneumonia. Eur. Respir. J. 2003;21: 713–719.

Maurin M, Raoult DF. Q fever. Am. Soc. Microbiol. 1999;12: 518–553.

McDade JE, Marrie TJ. Historical Aspects of Q Fever. In Q Fever: the disease. CRC Press, New York, USA; 1990. pp. 5–21.

Michelizzi VN, Dodson MV, Pan Z, Amaral MEJ, Michal JJ, McLean DJ, et al. Water buffalo genome science comes of age. Int. J. Biol. Sci. 2010;6: 333.

Nusinovici S, Madouasse A, Hoch T, Guatteo R, Beaudeau F. Evaluation of two PCR tests for Coxiella burnetii detection in dairy cattle farms using latent class analysis. PloS One. 2015;10: e0144608.

Paul S, Toft N, Agerholm JS, Christoffersen AB, Agger JF. Bayesian estimation of sensitivity and specificity of Coxiella burnetii antibody ELISA tests in bovine blood and milk. Prev. Vet. Med. 2013;109: 258–263.

Randhawa AS, Dhillon SS, Jolley WB. Serologic prevalence of Q fever in the state of Punjab, India. Am. J. Epidemiol. 1973;97: 131–134.

Raoult D, Marrie T. Q fever. Clin. Infect. Dis. 1995;489–495.

Roest HI, Bossers A, van Zijderveld FG, Rebel JM. Clinical microbiology of Coxiella burnetii and relevant aspects for the diagnosis and control of the zoonotic disease Q fever. Vet. Q. 2013;33: 148–160

Serrano-Pérez B, Almería S, Tutusaus J, Jado I, Anda P, Monleón E, Badiola J, et al. Coxiella burnetii total immunoglobulin G, phase I and phase II immunoglobulin G antibodies, and bacterial shedding in young dams in persistently infected dairy herds. J. Vet. Diagn. Invest. 2015;27: 167–176.

Shome R, Deka RP, Milesh L, Sahay S, Grace S, Lindahl JF. Coxiella seroprevalence and risk factors in large ruminants in Bihar and Assam, India. Acta tropica, 2019;194: 41–46.

Sodhi SS, Joshi DV, Sharma DR, Baxi KK. Seroprevalence of brucellosis and Q fever in dairy animals. Zbl. Vet. Med. B. 1980;27: 683–685.

Spiegelhalter DJ, Best NG, Carlin BP, Van Der Linde A. Bayesian measures of model complexity and fit. J. R. Stat. Soc. Series b (Stat Methodol). 2002;64: 583–639.

Spiegelhalter D, Thomas A, Best N, Lunn D. OpenBUGS user manual, version 3.0. 2. MRC Biostatistics Unit, Cambridge. 2007.

Sprong H, Tijsse-Klasen E, Langelaar M, De Bruin A, Fonville M, Gassner F, et al. Prevalence of Coxiella burnetii in ticks after a large outbreak of Q fever. Zoonoses Public Health. 2012;59: 69–75.

Toft N, Jørgensen E, Højsgaard S, Diagnosing diagnostic tests: evaluating the assumptions underlying the estimation of sensitivity and specificity in the absence of a gold standard. Prev. Vet. Med. 2005;68: 19–33.

Toft N., Innocent GT, Gettinby G, Reid SW. Assessing the convergence of Markov Chain Monte Carlo methods: an example from evaluation of diagnostic tests in absence of a gold standard. Prev. Vet. Med. 2007;79: 244–256.

Yadav JP, Malik SVS, Dhaka P, Kumar M, Bhoomika S, Gourkhede D, et al. Seasonal variation in occurrence of Coxiella burnetii infection in buffaloes slaughtered in India. Biol. Rhythm Res. 2019;1–7.

Vaidya VM, Malik SVS, Bhilegaonkar KN, Rathore RS, Kaur S, Barbuddhe SB. Prevalence of Q fever in domestic animals with reproductive disorders. Comp. Immunol. Microbiol. Infect. Dis. 2010;33: 307–321.

Waag DM. Coxiella burnetii: host and bacterial responses to infection. Vaccine, 2007;25: 7288–7295.

Walter SD, Irwig LM. Estimation of test error rates, disease prevalence and relative risk from misclassified data: a review. J. Clin. Epidemiol. 1988;41: 923–937.

Willems H, Thiele D, Frölich-Ritter R, Krauss H. Detection of Coxiella burnetii in cow’s milk using the polymerase chain reaction (PCR). J. Vet. Med. Series B. 1994;41: 580–587.

World Organisation for Animal Health (OIE) Terrestrial Manual. Chapter 3.1.16 Q Fever. 2018; Available from: https://www.oie.int/fileadmin/Home/eng/Health_standards/tahm/3.01.16_Q_FEVER.pdf

